# Novel epigenetic clock for fetal brain development predicts prenatal age for cellular stem cell models and derived neurons

**DOI:** 10.1101/2020.10.14.339093

**Authors:** Leonard C. Steg, Gemma L. Shireby, Jennifer Imm, Jonathan P. Davies, Alice Franklin, Robert Flynn, Seema C. Namboori, Akshay Bhinge, Aaron R. Jeffries, Joe Burrage, Grant W. A. Neilson, Emma M. Walker, Leo W. Perfect, Jack Price, Grainne McAlonan, Deepak P. Srivastava, Nicholas J. Bray, Emma L. Cope, Kimberly M. Jones, Nicholas D. Allen, Ehsan Pishva, Emma L. Dempster, Katie Lunnon, Jonathan Mill, Eilis Hannon

## Abstract

Induced pluripotent stem cells (iPSCs) and their differentiated neurons (iPSC-neurons) are a widely used cellular model in the research of the central nervous system. However, it is unknown how well they capture age-associated processes, particularly given that pluripotent cells are only present during the earliest stages of mammalian development. Epigenetic clocks utilize coordinated age-associated changes in DNA methylation to make predictions that correlate strongly with chronological age. It has been shown that the induction of pluripotency rejuvenates predicted epigenetic age. As existing clocks are not optimized for the study of brain development, we developed the fetal brain clock (FBC), a bespoke epigenetic clock trained in human prenatal brain samples in order to investigate more precisely the epigenetic age of iPSCs and iPSC-neurons. The FBC was tested in two independent validation cohorts across a total of 194 samples, confirming that the FBC outperforms other established epigenetic clocks in fetal brain cohorts. We applied the FBC to DNA methylation data from iPSCs and iPSC-derived neuronal precursor cells and neurons, finding that these cell types are epigenetically characterized as having an early fetal age. Furthermore, while differentiation from iPSCs to neurons significantly increases epigenetic age, iPSC-neurons are still predicted as being fetal. Together our findings reiterate the need to better understand the limitations of existing epigenetic clocks for answering biological research questions and highlight a limitation of iPSC-neurons as a cellular model of age-related diseases.

## Introduction

Induced pluripotent stem cells (iPSCs) offer a unique cellular system to investigate disease in human-derived cells. iPSCs are obtained by treating skin or blood cells with a set of core pluripotency transcription factors that reprogram the cells to a pluripotent state [1]. Established iPSC lines have the capacity to be further differentiated into specific cell types, including neurons, when treated with the appropriate factors [2–4]. This is of particular interest for neuroscience, as the only alternative cellular model for human neurons are immortalized cell lines. Because immortalized cell lines retain some physiological properties of the cancerous cells they were derived from [5] they do not fully recapitulate the neuronal phenotype. iPSC-derived neurons (iPSC-neurons), on the other hand, express appropriate morphological and neurophysiological properties of neurons and subject to different protocols can be differentiated into a wide range of specific neuronal subtypes [6]. iPSCs and their neuronal derivatives have been widely used to research disorders of the central nervous system, including developmental disorders such as autism and schizophrenia and age-related diseases such as Alzheimer’s disease (AD) and Parkinson’s disease. However, the extent to which iPSCs and especially iPSC-neurons capture age-associated processes is not known, which is fundamental to the study of age-related diseases. Of specific relevance is the fact that pluripotent cells only occur during the early stages of mammalian development and the effect of differentiation from iPSCs towards neurons on the developmental or aging trajectory of the cellular model [7] has yet to be adequately profiled.

Epigenetic mechanisms, such as DNA methylation (DNAm), are chemical processes that stably regulate gene expression, and while they are sensitive to environmental stimuli they also underpin key developmental processes [8–10]. There has been much interest and success in capitalizing on these patterns of epigenetic variation to derive individual age predictions from a biological sample. Age predictors based on DNAm, known as “epigenetic clocks” or “DNAm clocks”, are widely used to predict the “epigenetic age” of a sample. Epigenetic age, defined here as age predicted by an epigenetic clock, correlates strongly with chronological age, albeit not perfectly, and it has been hypothesized that the deviations from this prediction, referred to as age acceleration, are meaningful in the context of disease [11, 12]. The most well-known epigenetic clock is the Horvath multi-tissue clock (MTC) which was developed using a large number of samples (n > 8000) from 51 different tissues and cell types [13]. Overall, the MTC generates reliable predictions of chronological age for most sample-types, although there are potential biases when using Horvath’s clock in samples derived from certain tissues, especially the brain [14, 15]. To this end, a number of new DNAm clocks have been developed for specific tissue types, including whole blood [16] and cortex [14], which demonstrate more accurate predictions within the specified tissue. A less established refinement of epigenetics clocks is the application to specific developmental stages, with prenatal samples excluded or underrepresented in most training datasets. Recently, clocks were developed to predict gestational age (GA) of newborns, derived using pre- and perinatal DNAm data from blood samples [17] or placental samples [18]. While existing epigenetic clocks have been shown to accurately predict age in either postnatal brain samples (predominantly middle and older age) or non-brain prenatal samples, these tools have not been thoroughly tested on prenatal brain samples, and it is unknown whether they are able to delineate the earliest stages of brain development.

Previous analysis applying the MTC to DNAm data generated from iPSCs and their corresponding primary cells from adult donors found that the induction of pluripotency reversed the aging process, with iPSCs predicted as postnatal, with an epigenetic age close to zero (i.e. birth) [13]. As human pluripotent cells only occur during prenatal development, we hypothesize that existing clocks are not sensitive enough to accurately predict iPSCs at prenatal developmental stages. The inability to accurately estimate age during this crucial stage of neurodevelopment limits our ability to profile changes in epigenetic age induced by the differentiation of iPSCs into specific cell-types using already established DNAm clocks. Here we present a novel DNAm clock developed using prenatal brain samples that accurately predicts fetal age, outperforming other DNAm clocks in neurodevelopmental samples. We then apply our clock to iPSCs, iPSC-derived neuronal progenitor cells (NPCs) and iPSC-derived neurons, as well as in other cellular stem cell models and derived neuronal cells, to characterize the epigenetic age of these cellular models before and during the differentiation process.

## Methods

All statistical analyses were performed using R version 3.5.2 (https://www.r-project.org/) [19].

### Development of Fetal Brain Clock (FBC)

#### Description of fetal brain samples

To develop and profile the performance of the fetal brain clock (FBC), we collated a dataset of 258 fetal brain samples (see **Table S1**) of which 194 were processed by our group at the University of Exeter as described previously [20] and 64 were a subset (age < 0 years) of a publicly available dataset downloaded from the Gene Expression Omnibus (GEO; https://www.ncbi.nlm.nih.gov/geo/; GSE74193) [21]. Of the samples processed in Exeter, 154 overlap those included in [20] following additional outlier filtering by principal component analysis, where DNAm was quantified using the Illumina 450K DNA methylation array. The other 40 samples represent additional samples where DNAm was quantified using the Illumina EPIC DNA methylation array using a standard protocol as previously described [14].

#### Data pre-processing and quality control

All datasets for which raw data was available were pre-processed following a standard quality control (QC) and normalization pipeline as described before [14] using either the R package *wateRmelon* [22] or *bigmelon* [23]. Briefly, samples with low signal intensities or incomplete bisulfite conversion were excluded prior to applying the *pfilter*() function from the *wateRmelon* package, excluding samples with >1 % of probes with a detection P value >0.05 and probes with >1 % of samples with detection P value >0.05. This was followed by the exclusion of probes known to be affected by SNPs or known to cross-hybridize [24]. QC was finished by quantile normalization using the *dasen*() function of the packages *wateRmelon* or *bigmelon* [22, 23]. To harmonize the age variable across datasets, age was converted into days post-conception (dpc), as it represents the most precise unit of age available across the datasets. Where age was provided as weeks post-conception it was transformed to days post-conception by dividing by 7, and where age was reported in (negative) years it was transformed to days post-conception by multiplying by 365 and adding 280. Of note, a few samples (15 out of 258) are actually defined as embryonic (GA < 63 dpc) and not fetal.

#### Fetal brain clock development

To create two separate datasets for the purpose of training and testing the FBC, 75% of the samples from each dataset were randomly assigned into a training dataset (n = 193, age range = 37-184 dpc, age median = 99 dpc), while the remaining 25% included in the testing dataset (n = 65, age range = 23-153 dpc, age median = 99 dpc) (**Figure S1, Table S1**). There was no overlap in samples between the training and testing dataset. To simplify the FBC development, only probes available in all samples after QC were taken forward (n = 385,069 probes). To develop the fetal brain clock we applied an elastic net (EN) regression model, using the approach described by Horvath [13], regressing chronological age against DNAm level of all available probes. The EN algorithm selects a subset of DNA methylation probes that together produce the optimal prediction of the outcome, in this case chronological age, by combining ridge and LASSO (Least Absolute Shrinkage and Selection Operator) regression. Briefly, ridge regression penalizes the sum of squared coefficients while LASSO penalizes the sum of the absolute values of coefficients. EN is a combination of both methods, where the user specifies the extent of the mixing of the two methods as a number between 0 and 1, in our application this was set to 0.5 [25]. EN was implemented with the R package *GLMnet* [26]. The shrinkage parameter lambda was calculated using 10-fold cross-validation on the data, which resulted in a lambda of 3.27.

#### Statistical evaluation of FBC performance

To profile the performance of the FBC, we additionally tested three established DNAm clocks: Horvath’s multi-tissue clock (MTC) [13], Knight’s Gestational Age clock (GAC) [17] and Lee’s Control Placental epigenetic clock (CPC) [18]. The clocks were applied using the *agep()* function of the *wateRmelon* package [22], where the default estimates the MTC and other clocks (here the GAC and CPC) can be estimated by providing the necessary coefficient and intercept values. The predictive accuracy of each clock was profiled in each dataset by two measures: Pearson’s correlation coefficient with reported chronological age and root mean squared error (RMSE). To investigate potential effects of sex on the predicted epigenetic age, linear models were fitted in the testing and validation datasets with FBC predicted epigenetic age as dependent variable, chronological age and sex as main effects and an interaction of chronological age and sex.

### Validation of the Fetal Brain Clock (FBC) in additional fetal datasets and adult cortex

To further test the FBC, we used data from 96 additional fetal brain samples currently being assessed by our group (unpublished data), none of which overlapped with either the training data or test data described in the previous section, with DNAm quantified using the Illumina EPIC DNA methylation array. QC and normalization were performed as described above. We also included data from 33 fetal samples from two publicly available datasets on GEO (GSE116754 and GSE90871) [27, 28], where DNAm was quantified using the Illumina 450K DNA methylation array. Pre-processing and QC for the publicly available datasets was not performed in our lab as no raw data was available.

Age of all samples was converted to dpc as described above. The combined validation dataset has an age range of 42 – 280 dpc with the median at 112 dpc (**Figure S1, Table S1**). To evaluate the performance of the DNAm clocks in adult brain samples we utilized data from the Brains for Dementia Research (BDR) cohort previously generated by our group [14]. Briefly, these data consist of 1,221 samples from 632 donors (age range 41-104 years, median = 84 years), with DNA extracted from the prefrontal cortex (n = 610) and occipital cortex (n = 611). DNAm was quantified using the Illumina EPIC DNAm array, and were pre-processed using a standard QC pipeline as described in [14].

#### Statistical evaluation of FBC performance

The predictive accuracy of the FBC was profiled in each dataset by two measures: Pearson’s correlation coefficient with reported chronological age and root mean squared error (RMSE).

### Testing of Fetal Brain Clock (FBC) in cellular samples

#### iPSC - neuron samples

Five different DNAm datasets generated using iPSCs, iPSC-derived NPCs and iPSC-derived neurons were used to characterize epigenetic age of the neuronal cell model, details of which can be found in **Table S1.**

For two of these datasets (*Imm, Price*) DNAm data was generated by our lab in Exeter, where DNAm was quantified using the Illumina EPIC DNA methylation array. These were supplemented by three publicly available datasets, downloaded from GEO (*Nazor*, GSE31848, *Sultanov*, GSE105093, and *Fernández-Santiago*, GSE51921) consisting of Illumina 450K DNAm array data [3, 4, 29].

References describing the origin of cell lines and the different methods used for cell culture and differentiation are listed in **Table S1**. Pre-processing and QC for the *Nazor* and *Fernández-Santiago* datasets was not performed in our lab as no raw data was available.

#### iPSC – motor neuron samples

A dataset comprised of 23 cellular samples with two iPSC samples and 21 derived motor neurons was generated by our lab, with DNAm quantified using the Illumina EPIC DNAm array (**Table S1**). These data were QC’d following the pipeline described above.

#### ESC – neuron samples

Two publicly available cellular datasets with NPCs and neurons derived from embryonic stem cells (ESCs) were downloaded from GEO (*Nazor*, GSE31848, *Kim* GSE38214) [29, 30]. Both datasets consist of data quantified using the Illumina 450K DNAm array. Pre-processing and QC for both ESC – neuron datasets were not performed in our lab as no raw data was available.

#### Statistical comparison of cellular states

The FBC was applied to DNAm data for all cellular samples available. To test for differences in predicted epigenetic age between cell stages within each dataset, either two sample t-tests or ANOVA followed by Tukey HSD multiple comparison (when three cell stages were available), were used. To combine results across all iPSC - neuron datasets, a mixed effects linear model was fitted with predicted epigenetic age as the dependent variable, a fixed effect for cell stage represented as two dummy variables contrasting NPCs vs iPSCs and iPSC-neurons vs iPSCs as and a random effect (i.e. random intercept) for dataset.

## Results

### Fetal brain clock outperforms existing DNAm clocks at predicting age of prenatal brain samples

We applied EN regression to genome wide DNAm data from a subset of available prenatal brain samples (n = 193; **Table S1** and **Figure S1**) to develop the fetal brain clock (FBC). 107 DNAm probes were assigned non-zero coefficients and therefore were selected as the basis of the FBC (**Table S2**). We found no overlap in the DNAm sites selected for the FBC and DNAm sites used in the other established clocks tested in our analysis. Testing the FBC clock in an independent test dataset of fetal brain samples (**Table S1** and **Figure S1**) to evaluate its performance we found a strong linear relationship between chronological and predicted prenatal age (r = 0.80; **Figure 1A**) with the majority of samples predicted within 15 days of their actual chronological age (RMSE = 14.84 dpc). To benchmark the performance of our clock, we compared it to three existing DNAm clocks: Horvath’s MTC [13], Knight’s GAC [17] and Lee’s CPC [18]. These clocks were selected as they represent either the most well-established algorithm with the broadest applicability (MTC) or were specifically developed to predict pre- and perinatal gestational ages, albeit in non-brain tissue (GAC and CPC). Of note, the MTC only predicted 27 fetal brain samples (41.2%) as prenatal (dpc < 280) with a low correlation between chronological and predicted age (r_MTC_ = 0.06). This correlation is much lower than those reported in the original manuscript when Horvath tested the clock in adult samples [13], highlighting the challenges with extrapolating clocks to samples which were not well represented in model development. By comparison, the GAC and CPC perform better than the MTC, although they have smaller correlation coefficients (r_GAC_ = 0.52 and r_CPC_ = 0.76) and are associated with a larger error (RMSE_GAC_ = 21.32 and RMSE_CPC_ = 60.08) than the FBC. Interestingly, while the predictions from the GAC are more precise, it is not as effective at ranking the samples by age as the CPC. Taken together, these results demonstrate that our novel FBC outperforms existing clocks at predicting age in fetal brain samples, and therefore is the optimal tool available to profile the epigenetic age in models of neuronal development. When applying clocks to the training data, the three established clocks produce similar correlations and RMSEs as in the testing data. As expected, the predictions of the FBC in the training data are more accurate than the predictions in the testing data, reflecting overfitting of the model (**Figure S2**).

**Figure 1.**
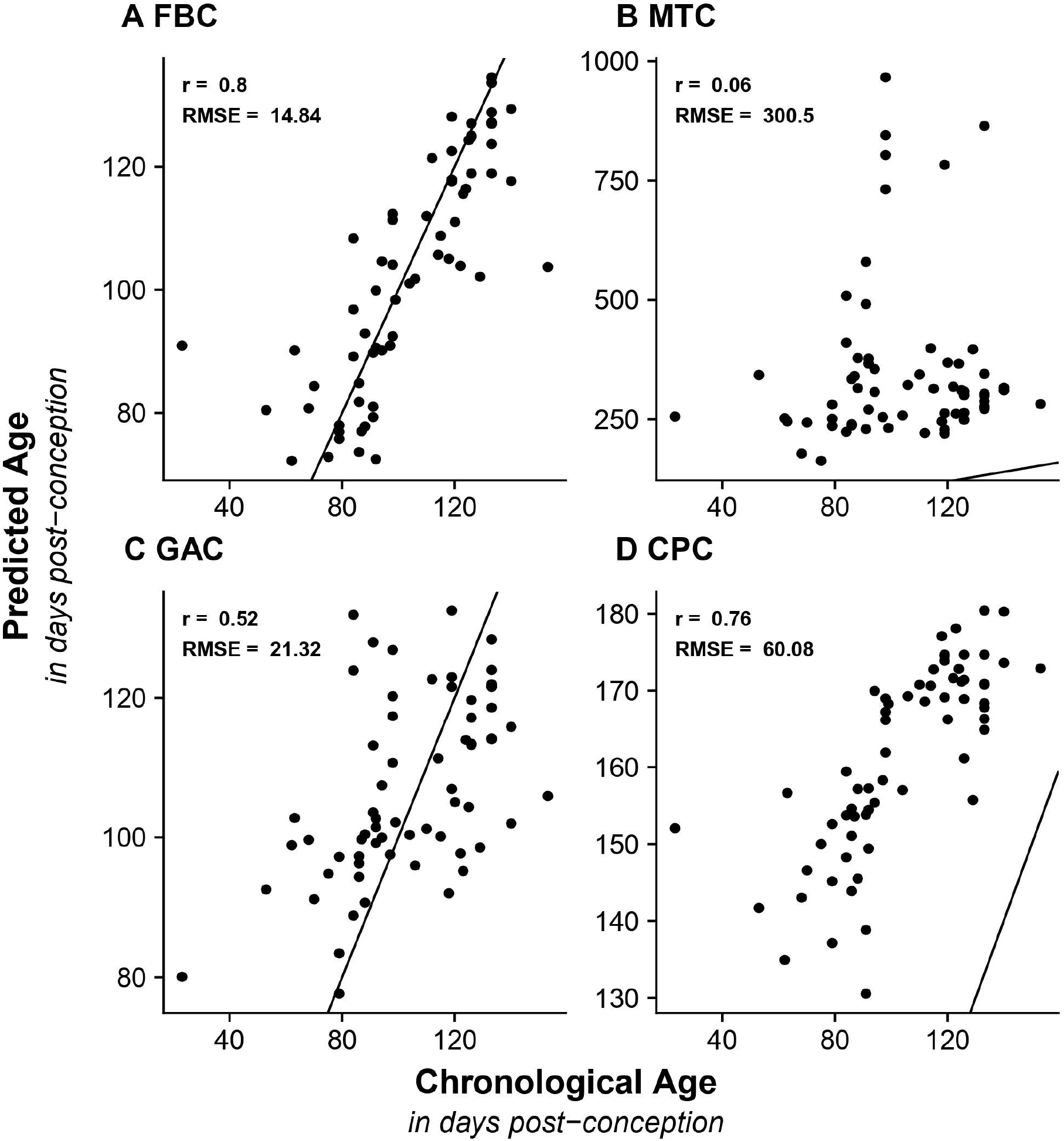
The Fetal Brain Clock (FBC) outperforms other DNAm clocks when applied to neurodevelopmental samples. Shown are scatterplots comparing chronological age (x-axis; days post-conception) against (y-axis; days post-conception) predicted epigenetic age calculated using **A** Fetal Brain Clock (FBC), **B** Horvath’s multi tissue clock (MTC), **C** Knight’s Gestational Age Clock (GAC), and **D** Lee’s Control Placental Clock (CPC) in our fetal brain testing dataset (n = 65, age range = 23 – 153 dpc). The black line indicates the identity line of chronological and predicted epigenetic age and represents a perfect prediction. Two statistics were calculated to evaluate the precision of each DNAm clock: Pearson’s correlation coefficient (r) and the root mean squared error (RMSE).

We further tested the FBC in an independent prenatal brain dataset (n = 129, **Table S1**), finding a stronger linear relationship between chronological age and predicted epigenetic age (r = 0.87, **Figure 2A**), than in the test dataset (**Figure 1A**) but a larger error rate (RMSE = 26.36 dpc). On closer inspection, we observed that this error is mainly driven by a subset of older samples (> 185dpc, **Figure 2A**) in the validation dataset (**Table S1**), that are older than any of the samples in the training data (**Figure 2**). If we limit our analysis to the samples whose chronological age overlaps the range of ages used in the training data (37 - 185 dpc; n = 125) then the error is decreased to 18.94 dpc. The performance of the established clocks in the validation dataset is also comparable to their performance in the testing dataset, with smaller correlations and higher error compared to the FBC and no sample with a predicted prenatal age by the MTC (**Figure 2B-D**).

**Figure 2.**
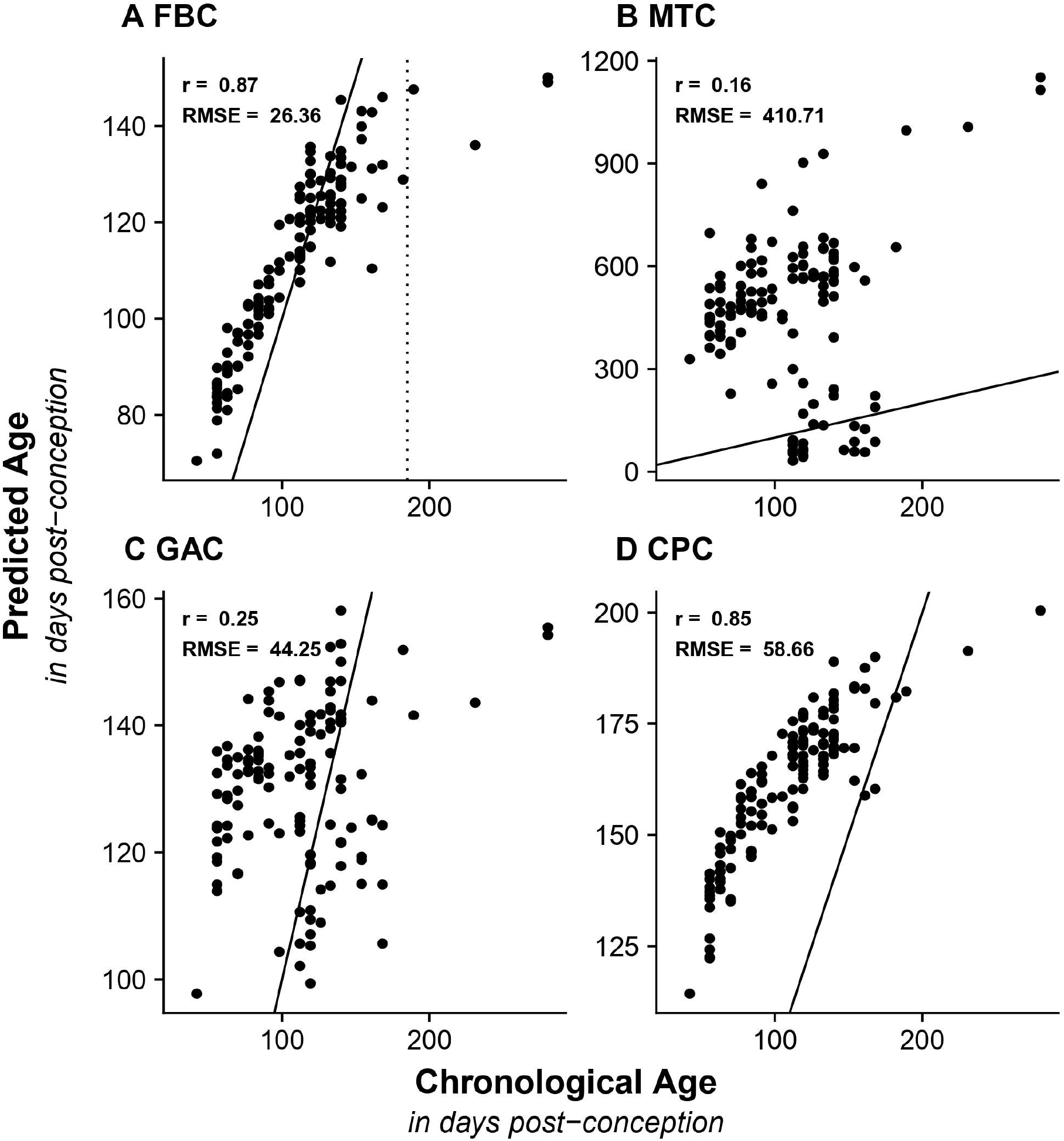
Validation of the Fetal Brain Clock in an independent fetal brain dataset. Shown are scatterplots comparing chronological age (x-axis; days post-conception (dpc)) against (y-axis; dpc) predicted epigenetic age calculated using **A** Fetal Brain Clock (FBC), **B** Horvath’s Multi Tissue Clock (MTC), **C** Knight’s Gestational Age Clock (GAC), and **D** Lee’s Control Placental Clock (CPC) on data from an independent validation dataset (n = 129, age range = 42 - 280). Two statistics were calculated to evaluate the precision of each DNAm clock: Pearson’s correlation coefficient (r) and the root mean squared error (RMSE). The dashed line in **A** indicates 185 dpc, which is the highest available age in the training dataset of the FBC.

Given our previous finding of divergent, sex-specific age trajectories at multiple DNAm sites during prenatal development [20], we tested whether the FBC performed differently between males and females in our testing dataset. Although this analysis initially indicated a significant difference in the correlation with age between males and females (P_Sex*Age_ = 0.0007), this relationship is likely driven by outliers. Indeed, a sensitivity analysis excluding the two samples with youngest and oldest predicted ages produced a non-significant result (P_Sex*Age_ = 0.081). Repeating the analysis in the validation dataset we find a small significant effect of sex on age in the full dataset (P_Sex*Age_ = 0.00179), which was driven by the samples older than 185, all of them being female. As samples older than 185 dpc produce inaccurate predictions, they could unfairly bias the analysis of potential sex effects and after removing the them from the analysis, there is no longer a significant effect of sex on the accuracy of the clock (P_Sex*Age_ = 0.95).

#### Fetal and gestational age clocks are not able to predict adult ages in adult brain tissue

All four clocks were additionally tested in an adult brain DNAm dataset (**Figure S3**). As expected, the FBC performs poorly in this sample set, with all samples predicted as prenatal albeit at the older end of the spectrum of ages in the training data (range of predicted ages 115-170 dpc). In contrast the MTC performs the best (r_MTC_ = 0.65, RMSE_MTC_ = 20.11 years) as it is the only clock we considered that was developed using adult samples. As with the FBC, the GAC and CPC fail to produce predictions of postnatal age, again reflecting the fact that they were also constructed using data from pre- or perinatal samples.

#### Fetal brain clock captures differences in differentiation of cellular stem cell models towards neurons

Having demonstrated that our novel FBC is the optimal clock to profile age in prenatal brain samples, we applied it to DNAm data from multiple cellular studies to characterize epigenetic age in iPSCs and ESCs differentiating towards cortical neurons. All samples were estimated to have a fetal epigenetic age, regardless of cell stage, cell line origin or differentiation protocol. Furthermore, they were predicted to have a “young” fetal age (**Figure 3**) with the iPSCs having a mean predicted age of 75.6 dpc (SD = 6.9 dpc; n = 59), the NPCs having a mean age of 79.1 dpc (SD = 11.0 dpc; n = 8) and iPSC - neurons having a mean predicted age of 83.2 dpc (SD = 8.87 dpc; n = 31). To test whether the differentiation process had an effect on the epigenetic age predictions from the FBC we compared the estimated ages between iPSCs and neurons, observing significant differences in all datasets (**Figure 3A**), with neurons being older than iPSCs. For the *Imm* dataset, which included proliferative NPCs as well as postmitotic neurons, we additionally found a significant difference between NPCs and iPSC-neurons (Δ_mean_ = 20.0 dpc, P = 0.00039), but not between iPSCs and NPCs (Δ_mean_ = 10.0 dpc, P = 0.24). In contrast, in the *Nazor* dataset, which only included iPSCs and NPCs, we did find a significant difference in predicted epigenetic age between iPSCs and NPCs (Δ_mean_ = 10.5 dpc, P = 0.00734). Meta-analyzing the data across the five studies including iPSCs and iPSC-derived NPCs and neurons, we found that iPSC-neurons were predicted to have a significantly advanced epigenetic age compared to iPSCs of about 2 weeks (Δ_mean_ = 13.35 dpc, P = 2.13e-11) but no significant difference was observed between iPSC and NPCs (Δ_mean_ = 4.33 dpc, P = 0.11).

**Figure 3.**
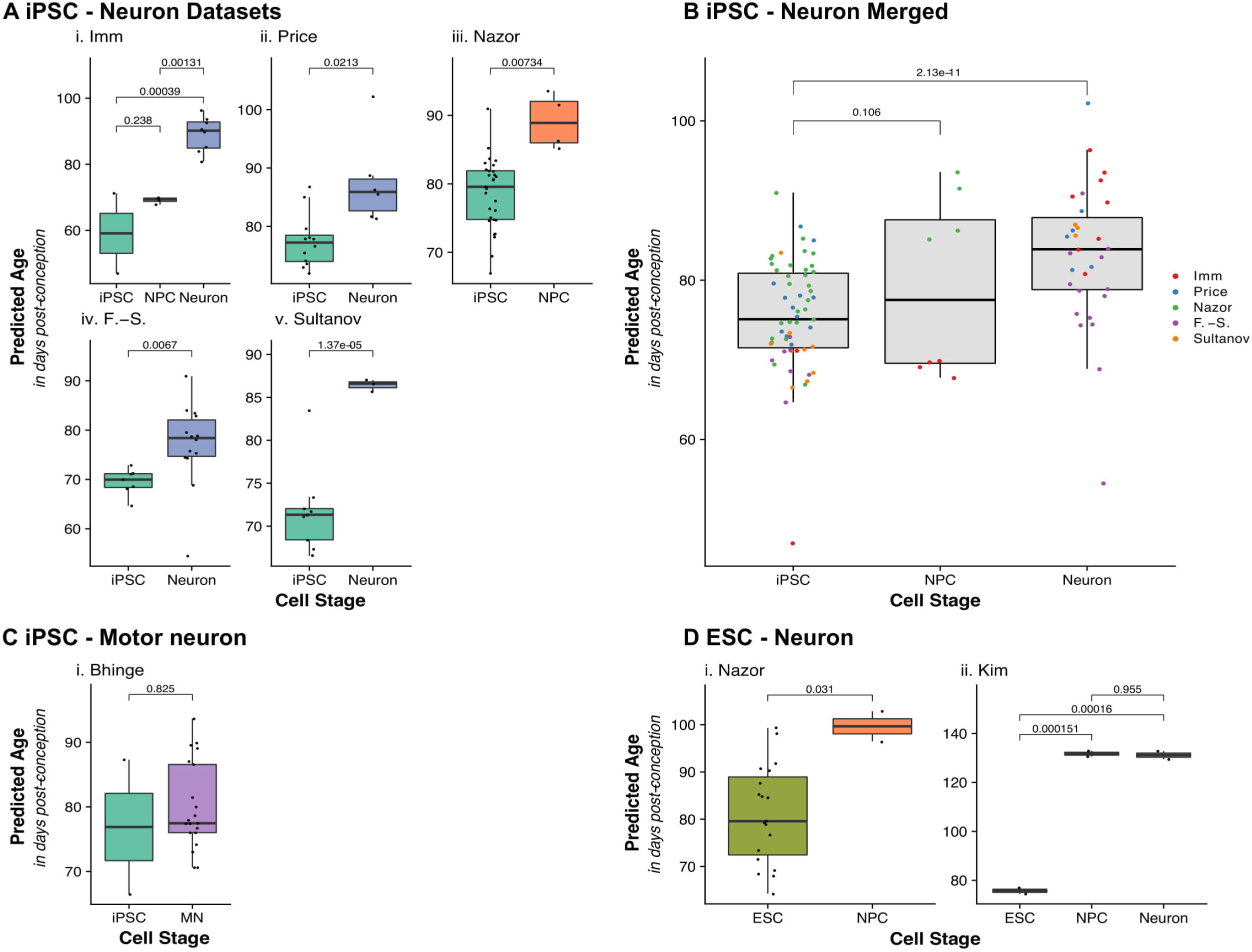
Comparisons of predicted epigenetic age using the fetal brain clock between cellular models throughout differentiation states. **A** Boxplots comparing the distribution of predicted epigenetic age (days post-conception) in iPSCs and their derived NPCs and neurons, where each panel represents a different dataset. P values of Tukey HSD corrected ANOVA for the *Imm* dataset and two-sample t-tests for *Price, Fernández-Santiago* and *Sultanov* datasets are given. F. -S. = *Fernández-Santiago*. **B** Boxplots of predicted epigenetic age calculated using the FBC where samples are grouped by cell stage (n = 82, 30 iPSCs, 4 NPCs, 48 iPSC-neurons) and colored by dataset. P values from mixed effects model are given for differences between iPSCs and NPCs (non-significant) and iPSC and neurons. **C** Boxplots comparing the predicted epigenetic age in a cohort with iPSCs and derived motor neurons. P values of two-sample t-test are given. **D** Boxplots of predicted epigenetic age by FBC applied on two datasets including ESCs and their derived NPCs and neurons. P values of Tukey HSD corrected ANOVA for the *Kim* dataset and two-sample t-tests for the *Nazor* dataset are given

As well as cortical neurons, we applied the FBC to DNAm data from a study on iPSCs and differentiated motor neurons. The motor neurons had a mean predicted epigenetic age of 79.82 dpc (SD = 6.82 dpc) which is slightly younger than the cortical neurons (mean_iPSC-neurons_ = 83.2 dpc). When comparing the predicted ages between the motor neurons and the iPSCs they were originally derived from, we did not observe a significant difference (Δ_mean_ = 2.94 dpc, P = 0.825, **Figure 3C**).

Finally, we tested for effects on epigenetic age through the process of differentiation from embryonic stem cells (ESCs) to NPCs and neurons, using two publicly available DNAm datasets (**Table S1**). In both datasets we observed a significant increase of predicted epigenetic age from ESC to NPC (**Figure 3D**). Additionally, in the *Kim* dataset, we were able to compare ESC derived neurons, and found a significant difference between ESCs and neurons (Δ_mean_ = 55.42 dpc, P = 0.00015, **Figure 3D**) but no change in epigenetic age from NPC to neuron (Δ_mean_ = -0.58 dpc, P = 0.95).

## Discussion

In this study we established a novel epigenetic clock, the fetal brain clock (FBC), to characterize the earliest stages of human neurodevelopment, and applied it to determine the epigenetic age of iPSCs and ESCs and their derived NPCs and neurons. Epigenetic clocks have been widely applied, including for the analysis of both in-vivo and in-vitro models of aging, where they have been shown to correlate with hallmarks of the aging process [31–34]. However, their application has predominantly been focused on studies involving adult samples. Given the lack of fetal brain samples in the development of existing DNAm clocks, prior to this study, there was no optimal method for estimating the age of fetal brain samples from DNAm data, limiting the ability to characterize iPSC-derived neuronal models or other models of neuronal development. We show that, in two non-overlapping independent validation datasets, the FBC generates predictions that correlate strongly with chronological age in prenatal brain samples. Furthermore, it outperforms both a pan-tissue epigenetic clock (Horvath’s MTC), and epigenetic clocks focused on the same developmental stage, but based on DNAm profiled in different tissues (Knight’s GAC and Lee’s CPC) [13, 17, 18]. The FBC outperforms these clocks using both correlation and error statistics (RMSE), indicating that it is not only better at ranking the samples, but it generates more precise estimates.

As the FBC was predominantly trained on second trimester brain samples, with some first trimester samples, it made less accurate predictions when applied to third trimester samples and performed extremely poorly in adult brain samples. Altogether, this reinforces the findings of previous studies that have also demonstrated that the applicability of DNAm clocks is dependent on the characteristics of the data are were trained on, with the tissue and age range of the training samples of particular relevance [14, 15]. More specifically, we note that while the accuracy of a DNAm clock is typically decreased in tissues not represented in its training data, clocks are completely limited to predicting ages represented in the training data. If the true age of a tested sample lies outside of the age range of the training data, the clock is unable to provide an appropriate prediction with the magnitude of inaccuracy increasing as the true age of the sample becomes more extreme suggesting that, in general, age range is more critical than tissue when training a clock.

Previous epigenetic clocks have shown that the predicted epigenetic age of iPSCs is significantly lower than the cells from which they are reprogrammed and the chronological age of the donor at sample donation [13]. The induction of pluripotency reprograms the epigenome, including at the loci used in the clock algorithm, ultimately leading to a younger predicted epigenetic age. However, in these analyses the predicted ages remain postnatal, which is unexpected as human pluripotent cells only occur during the early stages of human development and hence we hypothesized that, with an adequately calibrated clock, iPSCs would be expected to be estimated as being early fetal. Applying the FBC to five datasets of iPSCs and iPSC-derived NPCs and neurons, we found this to be the case. iPSCs were estimated as having a mean age of 75.6 dpc, fitting our hypothesis that they represent first trimester developmental stages. These results align with studies that have reported rejuvenation effects on the transcriptome, telomeres and mitochondria of iPSCs following reprogramming [35–37]. In addition, we profiled the effect on predicted epigenetic age following the differentiation of iPSCs towards neurons reporting a small but significant aging effect of 13 dpc. This developmental stage coincides with fetal neurogenesis [38] and suggests that while differentiation does induce an aging process, it does not accelerate iPSC-neurons to a postnatal state. Of note, Mertens and colleagues found that while iPSCs lose age related transcriptomic signatures, induced neurons (iNs; neurons directly reprogrammed from fibroblasts) keep their specific aging signatures [36]. Therefore, it would be interesting to apply our FBC to iNs, iPSCs, iPSC-neurons and their corresponding somatic tissues to verify whether age associated DNAm differences are also preserved in iNs. In addition, we tested the FBC on ESC and ESC-derived neurons. Although our sample size was small, the results also suggested that the differentiation process induced a small aging effect as the ESCs were differentiated into neurons. Altogether, our results indicate that iPSC-neurons may have limited utility for the study of age-related brain diseases, like Alzheimer’s disease or other dementias, as many molecular processes related to an aging phenotype may not be recapitulated.

Epigenetic clocks have been utilized for a wide range of applications [34], with a predominant focus on exploring the biological meaning of deviations between chronological age and epigenetic age. As we have shown the cause of this deviation may result from the use of an inappropriate clock, therefore the FBC is a critical tool for assessing whether epigenetic age acceleration during neurodevelopment is associated with later life outcomes such as disease or in utero exposures (such as maternal smoking). The FBC also has utility for determining the specific developmental stage a model of neurodevelopment recapitulates (e.g. brain organoids or cellular neuronal models) and how different exposures or genetic backgrounds may influence neurodevelopmental processes and aging.

While a strength of our study is the development of a bespoke clock to optimally profile the epigenetic age of human fetal brain samples, due to the training data predominantly containing second trimester samples the FBC is most accurate for this period of neurodevelopment. We are confident that the FBC has correctly predicted fetal epigenetic ages for the cell lines included in our analysis as the vast majority of the stem cell models and their derived neurons were less than the median age in the training data. This indicates that the predictions are not confounded by saturation of the coefficients. Although we took advantage of previously published data to include all available samples appropriate for addressing our research questions, some of the group sizes, in particular the NPCs, were small and we were not powered to detect significant aging effects as a result of differentiation from iPSCs to NPCs in all datasets. Furthermore, across the different studies there was variation in the estimated prenatal age of each cell state; we hypothesize that these differences results from subtle variation in differentiation protocols, timepoints of cell collection or the definition of NPCs within the respective studies [39]. Despite this, we are confident in the conclusions we report as these study specific effects were controlled for in our analysis.

In summary, we demonstrate that established DNAm clocks struggle to capture changes in epigenetic age during neurodevelopment and for precise predictions a bespoke clock trained on fetal brain data is required. Using the FBC to assess the epigenetic age of iPSCs and differentiated neurons, we found that iPSCs and derived NPCs and neurons reflect early prenatal developmental stages. Our findings question the suitability of the iPSC-neurons for the study of aging associated processes.

## List of abbreviations

Abbreviation: Definition
AD: Alzheimer’s Disease
BDR: Brains for Dementia Research
CPC: Control Placental Clock
DNAm: DNA methylation
dpc: Days post-conception
EN: Elastic net
ESC: Embryonic Stem Cell
FBC: Fetal brain clock
GA: Gestational age
GAC: Gestation age clock
GEO: Gene Expression Omnibus
iN: Induced neuron
iPSC: Induced pluripotent stem cell
iPSC-neuron: iPSC-derived neuron
LASSO: Least Absolute Shrinkage and Selection Operator
MTC: Multi-tissue clock
NPC: Neuronal precursor cell
QC: Quality control
RMSE: Root mean squared error

## Declarations

### Ethical approval

Ethical approval for the collection of human fetal brain tissue acquired from the Human Developmental Biology Resource was granted by the Royal Free Hospital research ethics committee under reference 08/H0712/34 and Human Tissue Authority material storage license 12220. Ethical approval for collection of human fetal brain tissue for the Medical Research Council Brain Bank was granted under reference 08/MRE09/38. Ethical approval for the work with methylomic data of human brain tissue from the BDR cohort was granted by the University of Exeter, College of Medicine and Health Research Ethics Committee under reference Mar20/D/009Δ5. Ethical approval for the Price cellular samples is as follows: participants were recruited and methods carried out in accordance to the ‘Patient iPSCs for Neurodevelopmental Disorders (PiNDs) study’ (REC No 13/LO/1218). Informed consent was obtained from all subjects for participation in the PiNDs study. Ethical approval for the PiNDs study was provided by the NHS Research Ethics Committee at the South London and Maudsley (SLaM) NHS R&D Office.

### Consent for publication

Not applicable

### Availability of data and materials

Raw and normalized DNAm data for the human fetal brain samples used to test and train the fetal brain clock have been submitted to the NCBI Gene Expression Omnibus (GEO; http://www.ncbi.nlm.nih.gov/geo/) under accession numbers GSE58885, GSE74193, GSE157908, GSE116754 and GSE90871. DNAm data for the unpublished fetal brain are available from the corresponding author on request. DNAm data for the cellular datasets, are also available under accession numbers GSE158089, GSE105093, GSE51921, GSE31848 and GSE38214. DNAm data for the cellular dataset *Bhinge* is available from the corresponding author on request. The coefficients of the FBC are included within the article in **Table S2**. The custom R scripts used for this study and to apply the FBC are available at https://github.com/LSteg/EpigeneticFetalClock.

## Authors’ contributions

L.C.S, G.L.S. and E.H. designed the work; J.M., E.H., E.L.D., N.J.B, A.B., J.P., D.P.S., N.D.A. and K.L. acquired funding; J.I., J.P.D., R.F., E.L.D., S.C.N., A.R.J., J.B., G.W.A.N., E.M.W., L.W.P., E.L.C., K.M.J., A.F., G.M., D.P.S., E.P., A.B., N.J.B. and N.D.A acquired and processed samples and data; L.C.S, G.L.S. and E.H analyzed and interpreted data; L.C.S. and E.H. wrote the paper.

## Acknowledgements

We would like to gratefully acknowledge all donors and their families for the tissue provided for this study. We are grateful to Rosy Watkins, Paulina Nowosiad, Rupert Faraway and Roland Nagy for generation and maintenance of *Price* iPSCs [41, 42].

## Additional Files

**Additional File 1: Table S1 Summary of fetal, adult and cellular datasets**.

**Additional File 2: Table S2 Intercept and coefficients of the fetal brain clock**.

**Additional File 3: Figure S1 Histogram of age distribution**. Chronological age of the fetal samples measured in days post-conception in A) training data, B) testing data and C) validation data.

**Additional File 4: Figure S2 Comparison of predictions from the four DNAm clocks in the training data (n = 193)**. Shown are scatterplots comparing chronological age (x-axis) against age (y-axis) predicted epigenetic age for **A** Fetal Brain Clock (FBC); **B** Horvath’s Multi Tissue Clock (MTC); **C** Knight’s Gestational Age Clock (GAC); **D** Lee’s Control Placental Clock (CPC) in the data used for training of the FBC. Where necessary, predicted age was converted to days post-conception. The black line indicates the identity line of chronological and predicted epigenetic age and represents a perfect prediction. Two statistics were calculated to evaluate the precision of each DNAm clock: Pearson’s correlation coefficient (r) and the root mean squared error (RMSE).

**Additional File 5: Figure S3 Comparison of predictions from the four DNAm clocks in adult brain samples (n = 1221)**. Shown are scatterplots comparing chronological age (x-axis) against (y-axis) predicted epigenetic age calculated using **A** Fetal Brain Clock (FBC); **B** Horvath’s multi tissue clock (MTC); **C** Knight’s Gestational Age Clock (GAC); **D** Lee’s Control Placental Clock (CPC) in an independent adult brain dataset. Where necessary, predicted age was converted to years, where 0 indicates birth. The black line indicates the identity line of chronological and predicted epigenetic age and represents a perfect prediction. Two statistics were calculated to evaluate the precision of each DNAm clock: Pearson’s correlation coefficient (r) and the root mean squared error (RMSE).

