## Additional File 1 Table S1 for "Novel epigenetic clock for fetal brain development predicts prenatal age for cellular stem cell models and derived neurons"

| **Dataset** | **Cohort / Study** | **BeadCHIP Array** | **N** | **Sex** | | | **Age** (in days post-conception) | | | **References** | |
| --- | --- | --- | --- | --- | --- | --- | --- | --- | --- | --- | --- |
|  |  |  |  | **Female** | **Male** | **NA** | **Median** | **Range** | |  | |
| **Fetal** | Training | 450K / EPIC | 193 | 83 | 110 |  | 99 | 37 | 184 | [20, 21] | |
|  | Testing | 450K / EPIC | 65 | 34 | 31 |  | 99 | 23 | 153 | [20, 21] | |
|  | Validation | 450K / EPIC | 129 | 61 | 58 |  | 112 | 42 | 280 | [27, 28] | |
|  |  |  |  |  |  |  | **Age** (in years) | | |  | |
|  |  |  |  |  |  |  | **Median** | **Range** | **Median** |  | |
| **Adult** | BDR | EPIC | 1221 | 577 | 644 |  | 84 | 41 | 104 | [14] | |
|  |  |  |  |  |  |  | **Cell Stage** | | | **Cell Line References** | **Differentiation Protocol Reference** |
|  |  |  |  |  |  |  | **iPSC** | **NPC** | **Neuron** |  |  |
| **iPSC - Neuron** | Imm | EPIC | 14 | 14 |  |  | 2 | 4 | 8 | [39] | [2] |
|  | Price | EPIC | 18 | 2 | 15 | 1 | 12 |  | 6 | [41] | [42] |
|  | Nazor | 450K | 33 | 29 | 4 |  | 29 | 4 |  | [29] | [29] |
|  | Fernández-Santiago | 450K | 21 | 10 | 11 |  | 7 |  | 14 | [3] | [3] |
|  | Sultanov | 450K | 12 |  |  | 12 | 9 |  | 3 | [4] | [4] |
| **iPSC - MN** | Bhinge | EPIC | 23 |  |  | 23 | 2 |  | 21 | [40] | [40] |
|  |  |  |  |  |  |  | **Cell Stage** | | |  |  |
|  |  |  |  |  |  |  | **ESC** | **NPC** | **Neuron** |  |  |
| **ESC - Neuron** | Kim | 450K | 6 | 6 |  |  | 2 | 2 | 2 | [30] | [30] |
|  | Nazor | 450K | 21 | 13 | 8 |  | 19 | 2 |  | [29] | [29] |
