## Additional File 2 Table S2 for "Novel epigenetic clock for fetal brain development predicts prenatal age for cellular stem cell models and derived neurons"

| **Name** | **Coefficient** |
| --- | --- |
| (Intercept) | -58.24134133 |
| cg00187437 | 16.27245218 |
| cg00282195 | 1.269269438 |
| cg00506250 | -5.750181404 |
| cg01081189 | -22.78182971 |
| cg01368219 | 39.92951171 |
| cg01395533 | 1.691597392 |
| cg01502919 | -1.174000173 |
| cg01568521 | 51.48749285 |
| cg01613337 | 14.83607261 |
| cg01734786 | -4.145713875 |
| cg01907457 | 0.748167644 |
| cg02060096 | -11.30081786 |
| cg02372820 | 1.196850196 |
| cg02905964 | -2.404271849 |
| cg03037561 | 1.000469483 |
| cg03226872 | -0.257371533 |
| cg03549739 | 8.077886836 |
| cg03870397 | -9.171149711 |
| cg04555379 | 14.02551628 |
| cg04581294 | 2.286808978 |
| cg04657146 | -4.467863354 |
| cg04831806 | -2.61536497 |
| cg05401839 | 1.469960345 |
| cg05785348 | -0.94813745 |
| cg05991401 | -1.842631263 |
| cg06301399 | 30.50647565 |
| cg06341590 | -7.753793933 |
| cg06413196 | -0.38682967 |
| cg06480764 | -26.93920878 |
| cg06532546 | 6.822472124 |
| cg06662568 | 21.21102654 |
| cg06683316 | 9.84075194 |
| cg07168142 | 3.248573684 |
| cg07408340 | 9.731048121 |
| cg07548255 | -0.198339056 |
| cg08331635 | -9.957767545 |
| cg08349573 | -18.18049772 |
| cg08607533 | -5.419018045 |
| cg09179079 | -1.574959207 |
| cg09254686 | -2.778030845 |
| cg09275655 | -11.43992654 |
| cg10061292 | -30.99894233 |
| cg10266681 | -2.250263683 |
| cg10280963 | 10.20006115 |
| cg10976968 | -1.749312271 |
| cg10981717 | -6.614825156 |
| cg11067714 | 13.99743978 |
| cg11205312 | -9.421989326 |
| cg11315449 | 16.45657707 |
| cg11398400 | 8.632416158 |
| cg11531272 | 0.97038092 |
| cg12158056 | -10.81226478 |
| cg12315892 | -6.809567874 |
| cg12641286 | 3.611296377 |
| cg12809031 | -0.612836722 |
| cg13617904 | -4.311034033 |
| cg13861904 | -0.52301438 |
| cg13861948 | 21.0941737 |
| cg14112569 | 4.184280401 |
| cg14198472 | -6.913673506 |
| cg14243741 | 2.388057511 |
| cg14421348 | -21.92232107 |
| cg14996263 | -0.302240592 |
| cg15637415 | 11.61724362 |
| cg15673034 | 8.727390653 |
| cg15770585 | -1.449612346 |
| cg15776560 | 30.33962208 |
| cg16586538 | -7.856695034 |
| cg16807679 | 3.144583019 |
| cg16918684 | 5.512977175 |
| cg17098103 | -3.65434643 |
| cg17540159 | 3.729137119 |
| cg17754515 | -26.74788549 |
| cg17839314 | -11.60106235 |
| cg17998566 | -21.22080046 |
| cg18645906 | 22.09466603 |
| cg19354746 | -1.364409926 |
| cg19451168 | 0.661939777 |
| cg19612309 | 4.761847353 |
| cg19927885 | 2.092488705 |
| cg20039211 | -3.75915027 |
| cg20282814 | 18.93004188 |
| cg20634498 | 1.633662784 |
| cg20660624 | 3.387472517 |
| cg20752831 | -3.646357808 |
| cg20778582 | -2.594477482 |
| cg20917891 | -13.08420368 |
| cg21554895 | 5.409038529 |
| cg21748136 | 0.619232136 |
| cg21859594 | 12.86716939 |
| cg22363368 | -0.749509176 |
| cg22764289 | -1.842314387 |
| cg22865215 | 60.2468917 |
| cg22891413 | 1.2084383 |
| cg23256459 | 2.577214702 |
| cg23437973 | -31.29787548 |
| cg23731272 | 4.657050639 |
| cg24129977 | -1.06778635 |
| cg24351977 | 2.290766621 |
| cg25309759 | -11.17449467 |
| cg25451082 | -19.75511753 |
| cg25581090 | 0.557856724 |
| cg25918267 | -6.031826965 |
| cg26450801 | 0.349103356 |
| cg26666107 | 5.973259312 |
| cg27455098 | -16.31062985 |
| cg27640020 | 0.061136687 |
