## Supplementary figures and images for "Novel epigenetic clock for fetal brain development predicts prenatal age for cellular stem cell models and derived neurons"

### Additional File 3 Figure S1

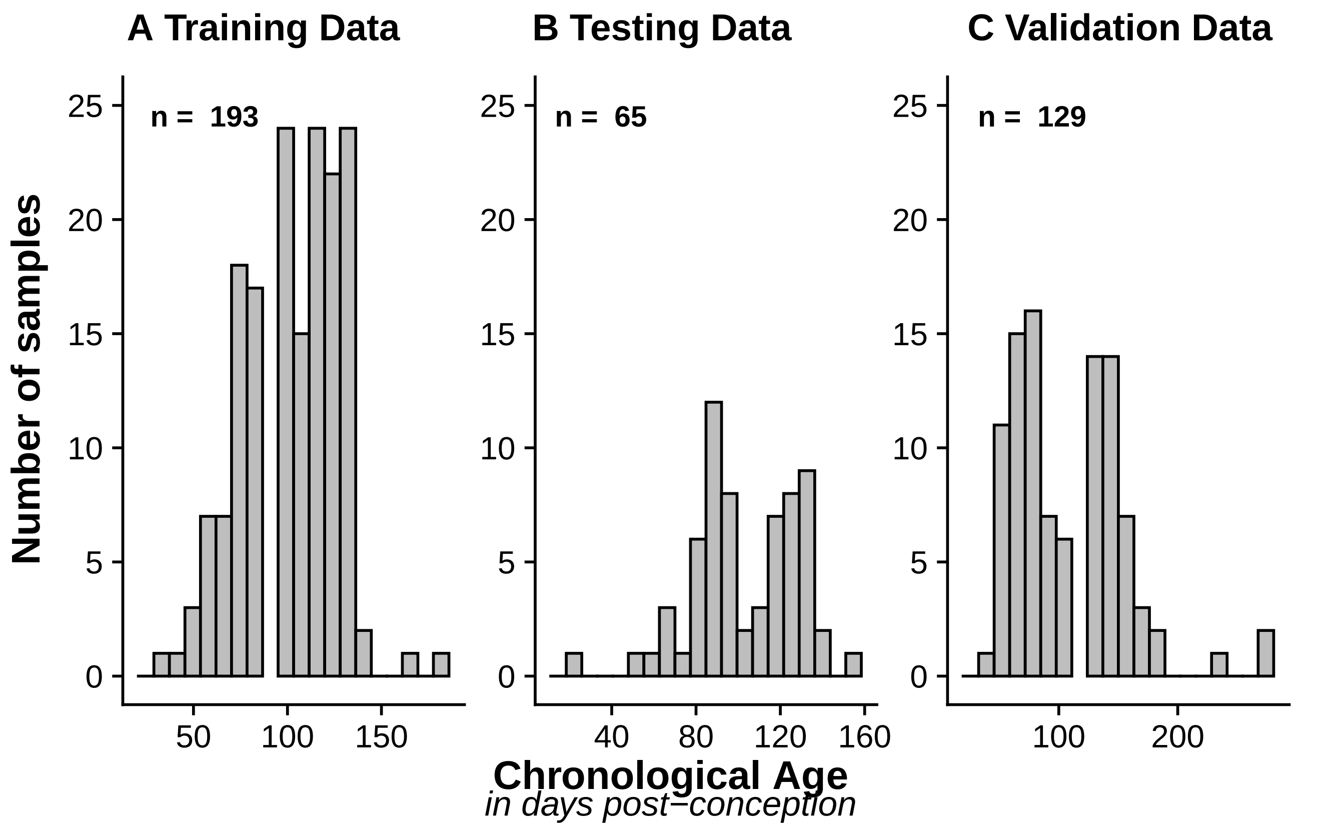

### Additional File 4 Figure S2

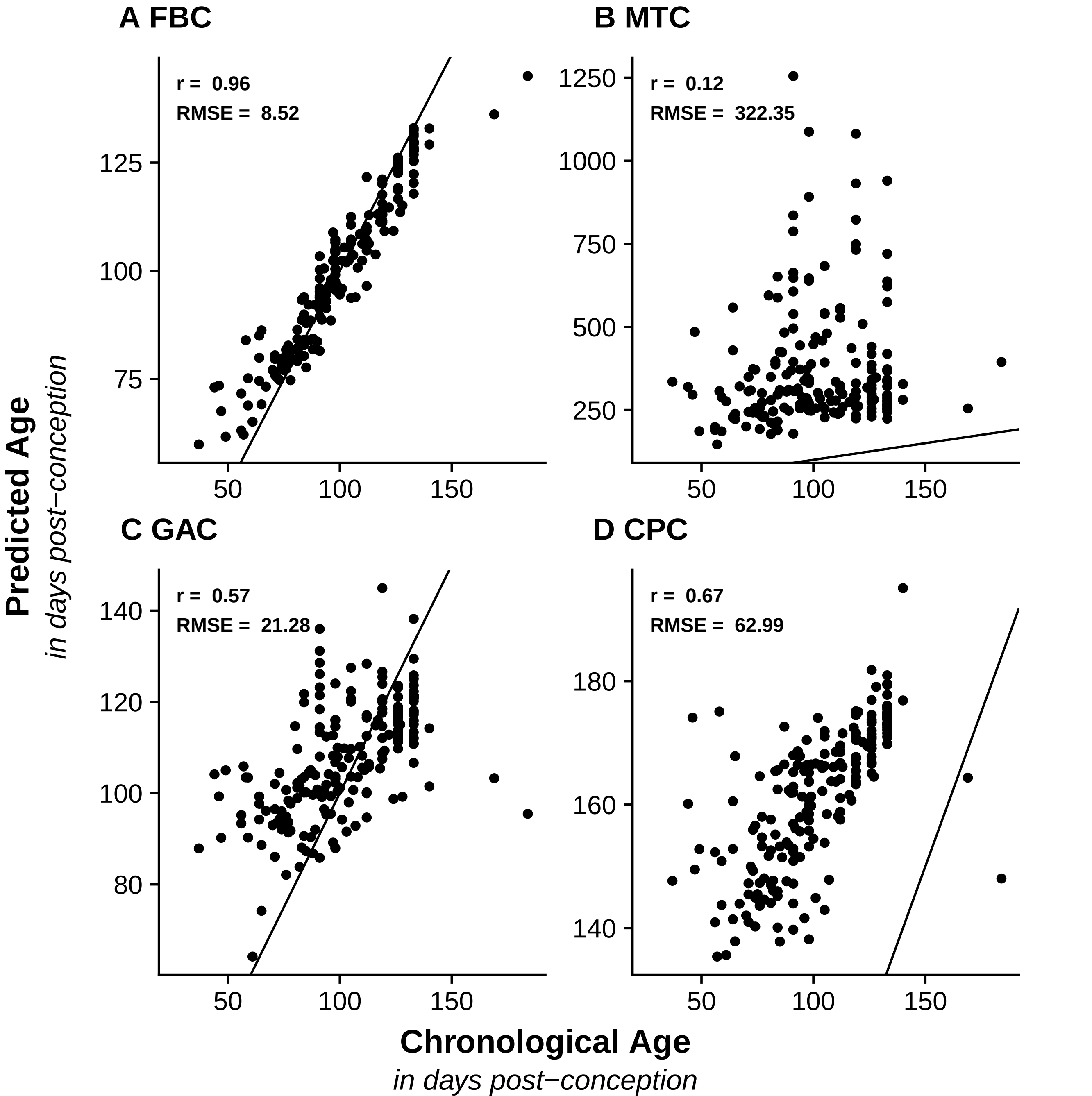

### Additional File 5 Figure S3

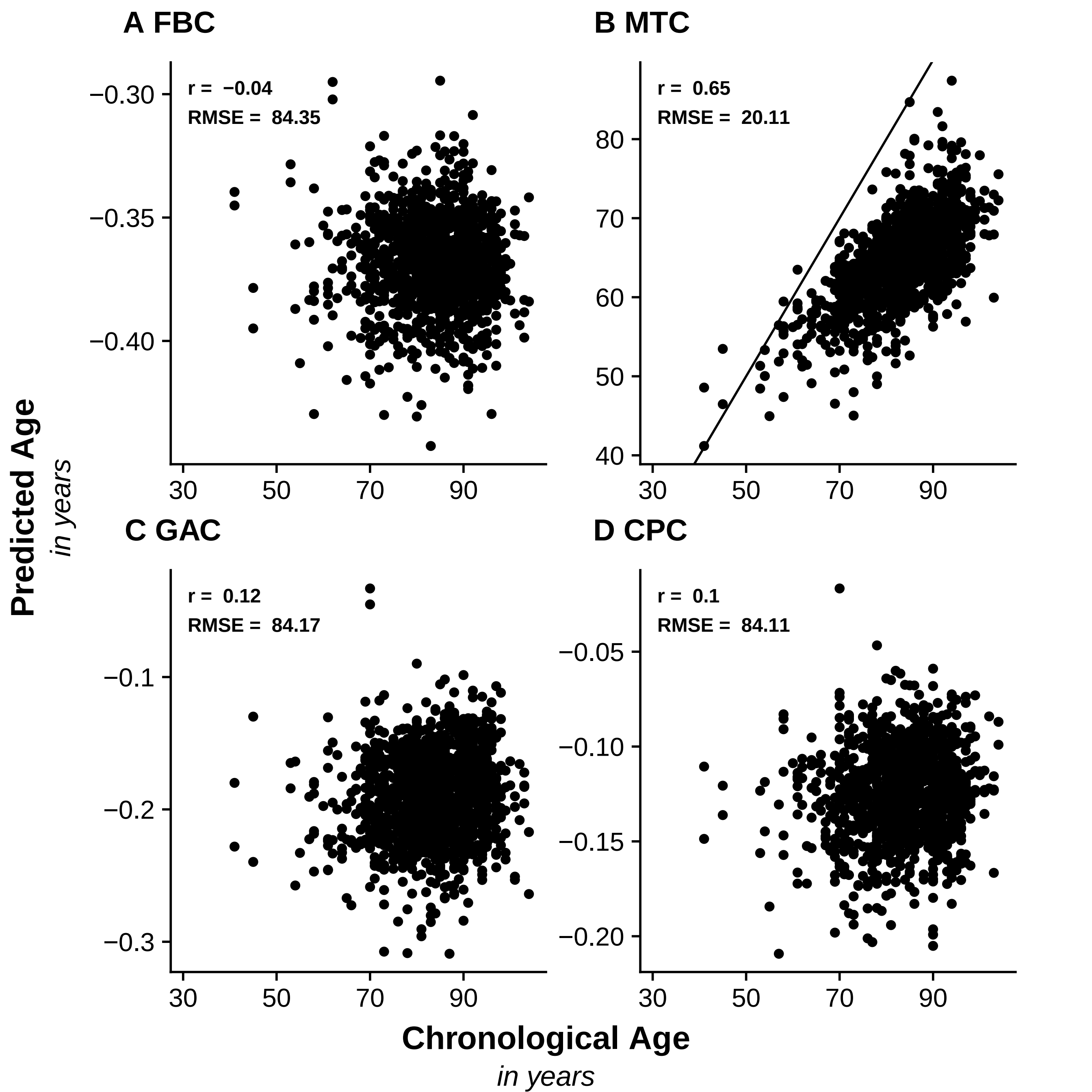
